# Neutral speciation in realistic populations

**DOI:** 10.1101/2024.01.10.575124

**Authors:** Erik D. Nelson

## Abstract

de Aguiar et al. have shown that basic patterns of species diversity found in nature can be described by a neutral model of speciation in which species emerge simply as a consequence of local mating, and mate preference for genetic similarity. Their results have been cited as support for the neutral theory of biodiversity. However, because the mutation rates considered in their work are much larger than those experienced by living organisms, there is still some question as to whether speciation will occur in this type of model under realistic conditions. Here, I develop a variant of the neutral model that includes a realistic mechanism for organism dispersal. I explore speciation in the model for a class of mobile organisms (butterflies), and I find that speciation does occur under conditions consistent with butterfly populations, albeit on narrow landscapes. The model also appears to exhibit scaling behavior – specifically, if the model is “scaled up” by increasing the area of the landscape while holding its length to width ratio and population density constant, the number of species tends to an asymptotic value. The results suggest that it is possible to infer speciation patterns in large populations by simulating much smaller, computationally tractable populations.

## Introduction

A classical view of speciation is that reproductive barriers between species emerge as a consequence of genetic divergence. In other words, genetically different individuals in a population are less likely to mate, or when they do mate, are more likely to produce offspring of lower fitness. Under these conditions, a population may progressively divide into sub–populations, for example, through genetic drift and isolation by distance, that are rarely able to form viable hybrids via interbreeding. This sort of reasoning has led to models of speciation in which mate preference or offspring fitness depend explicitly on the number of genetic differences between mating partners [1–4]. These types of models are often described collectively as extensions of the Dobzhansky–Muller model (for an excellent review of these models see [5, 6]).

In recent work, de Aguiar et al. have investigated a series of speciation models based on this principle [7–14]. Their approach is a extension of the random mating model developed by Higgs and Derrida to local mating in spatially distributed populations. In both versions of the model, genomes are described as sequences of binary “genes”; Mutations are neutral, and occur at a rate *µ* per gene per generation. Individuals *α* and *β* are permitted to mate only when *q*^*αβ*^, a measure of genetic similarity, exceeds a threshold value, *q*_*min*_. In the local mating version of the model, the potential mates of a focal individual are limited to a specific range of its position on a two–dimensional landscape. Under certain conditions (e.g. on population density, mutation rate, etc.) de Aquiar et al. find that a genetically uniform population will, over time, spontaneously divide into reproductively isolated clusters of individuals – i.e., into species – a process they refer to as “topopatric” speciation. While it is thought that speciation is usually driven by natural selection [15], their approach is remarkably successful in predicting the basic patterns of biodiversity found in nature [7,12]. However, because the mutation rates considered in their work are much larger than those experienced by real organisms, it is still unclear whether topopatric speciation will occur under realistic conditions.

To explain this, it is necessary to be specific about what a gene in the model represents in an organism. As an example in this work, I focus on butterfly populations. In order to estimate the model mutation rate *µ* for butterfles, I use the data obtained for Heliconius, an intensively studied genus of butterfly [16, 17]. Heliconius is a well known example of hybrid and ecological speciation, and is therefore an unlikely candidate for description by the neutral model (see however [14]). However, the data for Heliconius is accurate, and should provide reasonably good estimates for other species. The most recent estimate for the mutation rate in Heliconius is 2.9 *×* 10^−9^ per nucleotide site per generation [18].

The effective population size corresponding to this estimate is *N* = 2 *×* 10^6^. The average length of a Heliconius gene can be estimated from Drosophila (an organism used to identify butterfly genes) as Δ*L* ⋍ 1770 nucleotides [19], which results in a mutation rate per gene of about *µ* = 5.1*×*10^−6^ per generation. If we assume that a mutation in the neutral model corresponds to a single nucleotide mutation in a Heliconius gene, then the scaled mutation rate *Nµ* ⋍ 10 is well above the minimal value *Nµ* = 0.025 that leads to speciation in [7]. In random mating models such as the Wright–Fisher model [20], populations are known to obey certain “scaling” rules [21] which describe how a model will react when it is scaled up to larger population sizes. For example, under pure genetic drift, populations with the same values of *Nµ* but different values of *N* and *µ* have similar equilibrium distributions. Since the scaled mutation rate for Heliconius satisfies *Nµ >* 0.025, it is tempting to think that speciation will occur in the model for similar values of *N* and *µ*.

However, there are a number of problems with this analysis. First, there is no guarantee that the scaling rules [21] which hold for random mating models [22] will hold for populations with local mating on spatially extended landscapes. Assuming they do hold, the analysis does not account for empirical genetic differences between incipient butterfly species, which are typically on the order of 1 percent or more [16, 17, 23]. As a result, orthologous genes sampled from different species will differ at many more nucleotide sites than allowed by the condition assumed above.

This problem can be addressed by reconsidering the definition of a model gene. Let *d*^*αβ*^ denote the genetic (Hamming) distance between genomes *g*^*α*^ and *g*^*β*^, and, for simplicity, assume that distances between diploid genomes are measured by comparing paternal chromosomes only (i.e., so that *d*^*αβ*^ corresponds to the number of sites that differ between paternal chromosomes). The degree of similarity between model genomes *g*^*α*^ and *g*^*β*^ can be defined as

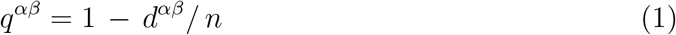

where *n* is the number of genes in a model genome. The fractional difference between incipient butterfly species is then *f* = *d*^*αβ*^*/ L*, where *L* is the length of sequence compared. In this work, I will consider that model genes refer only to segments of an organismal genome that potentially affect reproductive isolation. As noted above, mating between individuals *α* and *β* in the model can occur only when *q*^*αβ*^ is greater than some *q*_*min*_. If, at the same time, we require genetic differences between butterfly species to be at least 1 percent, and we again assume that a mutation in the model corresponds to a single nucleotide mutation in a butterfly gene, we obtain the following correspondence condition between genes in the model and genes in incipient species

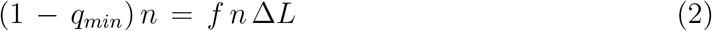

or simply Δ*L* = (1 − *q*_*min*_)*/f*, where Δ*L* is the organismal sequence length (to be deter-mined) represented by a model gene, and *f* = 0.01. In other words, given that a mutation in the model corresponds to a single nucleotide mutation in an organismal ”gene” (of length Δ*L*), the threshold number of mutational differences between model species is equal to the number of mutational differences between incipient butterfly species. Let *d*_*max*_ denote the value of *d*^*αβ*^ in Eq. (1) for which *q*^*αβ*^ satisfies *q*^*αβ*^ = *q*_*min*_ (here, *d*_*max*_ corresponds to the parameter *G* in [7]). The value of *d*_*max*_ used by de Aguiar et al. is approximately *d*_*max*_ = 0.06*n*, leading to a similarity threshold *q*_*min*_ = 0.94 and a gene length Δ*L* = 6 nucleotides. Clearly, Δ*L* no longer represents a gene in the literal sense, however, as this language is inherited from the model [8], it is convenient to continue using it below. Finally, using the data above for Heliconius, one obtains *µ* ⋍ 1.7 *×* 10^−8^ per gene per generation (about three orders of magnitude smaller than the minimal value of *µ* used by de Aguiar et al.) and *Nµ* ⋍ 0.035. Thus, according to these rough calculations, *Nµ* still satisfies the condition for speciation noted above.

There are a number of qualifying remarks to be made about this estimate. First, while I have used the effective population size for Heliconius to determine *Nµ*, it may be more accurate to consider the actual (census) population size, which is usually larger. In fact, even the effective population size for Heliconius populations can reach numbers as large as 10^7^ individuals [17]. At the same time, while I have used the background mutation rate for Heliconius to estimate *Nµ*, not all mutations to organismal “genes” will affect reproductive isolation. Finally, population sizes and nucleotide mutation rates vary among organisms (mammals, fish, birds, etc. [24]). While it is of interest to explore all of these conditions, simulations of the neutral model for realistic populations (*N >* 10^5^) are daunting, even on modern parallel computers (see below). In addition, data for many different organisms [24] lead to values of *Nµ* that are similar in order of magnitude to the value above estimated using Heliconius data given similar values for *f* and *d*_*max*_. Thus, simulations of the neutral model under conditions for butterfly populations should shed some light on whether topopatric speciation occurs in general. Does topopatric speciation occur under these conditions?

In this work, I develop a straightforward version of the neutral model that incorporates the flight patterns of butterflies in order to address this question. Individuals in the model have diploid genomes, and evolve on a continuous landscape. Mating is sexual, and occurs in discrete generations. In each generation, each female in the population randomly selects a mating partner from among the male individuals within a radius *R* of its current position. The mean number of offspring produced by each female is determined so as to maintain the local density of individuals at a fixed mean value. Offspring then disperse from the mating site in random directions according to an exponential flight length distribution [25] to start the next generation.

I explore the behavior of the model as it is “scaled up” to realistic population sizes and mutation rates by increasing the area of the landscape at constant population density while holding *Nµ* fixed at a value consistent with Heliconius data. Due to the large population sizes considered here, I employ a novel approach to define species clusters. Instead of solving the cluster problem for the whole population [9, 14], I sample a fraction of the population, and use a rapid clustering algorithm [26,27] to partition the sampled genomes by genetic distance. The clustering algorithm iteratively divides a sample into increasing numbers of clusters, and I select the partitioning with the largest number of clusters subject to the condition that the fixation index, *F*, a widely used measure of distance between genome clusters, satisfies *F > F*^⋆^ for all pairs of clusters in the partitioning, where *F*^⋆^ ⪞ 0.25, consistent with incipient species in natural populations [16,17,28] (later below, I demonstrate that this condition leads to very reasonable results for within species and between species genetic distances). The general behavior of the model as a function of population density, population size, and time is qualitatively similar to the models studied by de Aguiar et al. However, I find that persistent species occur only on narrow landscapes as realistic values of *N* and *µ* are approached. In this limit, species numbers for populations with the same density, on landscapes with the same length to width ratio scale with *Nµ* as noted above. Below, I describe the model in more detail, and then discuss the results of my simulations.

## Model

The genetic component of the model is similar to that considered by Schneider et al. [11]. Model genomes consist of two binary chromosomes, each of length *n* genes. During mating, genomes undergo a model form of meiosis (Fig. 1). In male meiosis, each chromosome is duplicated and crossovers occur at random between maternal and paternal copies at a rate of 1 crossover per meiosis resulting in a set of four model gametes [29]. The same procedure is conducted for female genomes but with no crossing over, consistent with butterfly reproduction. A gamete is selected at random from each set, and the selected gametes are combined to form a given offspring genome. The sex of an offspring is assigned at random. This process (including meiosis) is repeated a number of times for each mating event, depending on the number of offspring to be produced for a given female (see below).

**Fig. 1.**
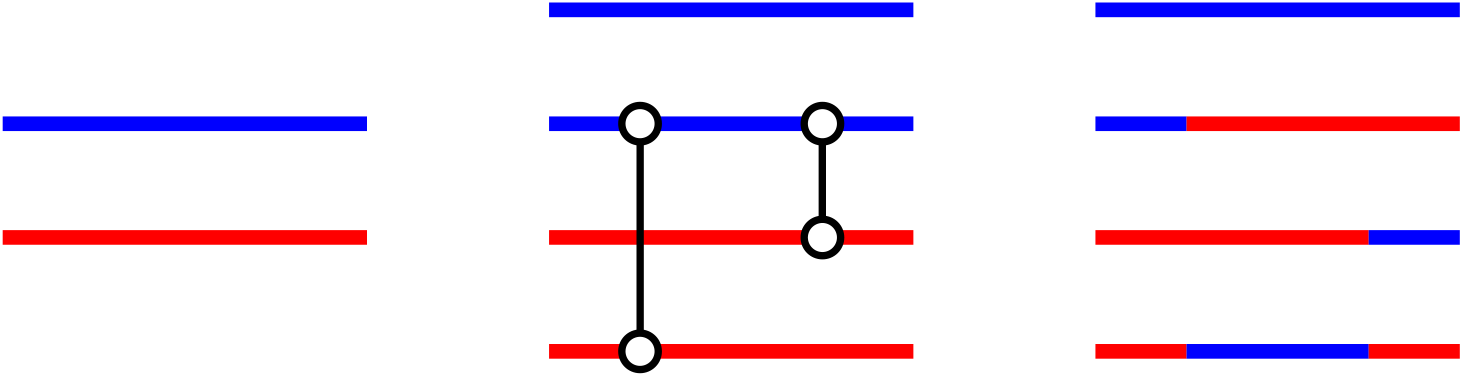
Model meiosis. Chromosomes (left) are duplicated to produce a set of four model chromatids (middle). Random connections (vertical lines) occur between maternal (red) and paternal (blue) chromatids, and genetic material is exchanged to the right of the connection points (circles) producing a set of four model gametes (right).

Genetic distances between genomes are determined according to the “incomplete dom-inance” scheme used by Schneider et al. [11]: Let 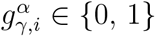 denote the state of gene *I* on chromosome *γ* ∈ {1, 2} in genome *α*, and let 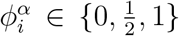 denote the average of 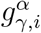 over 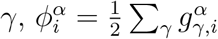. Distances between genomes are defined as,

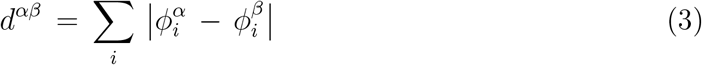

such that the effects of mutations at site *i* in chromosomes *γ* ∈ {1, 2} are additive. The fixation index for a pair of genome clusters *µ* and *ν* ≠ *µ* is computed as [30]

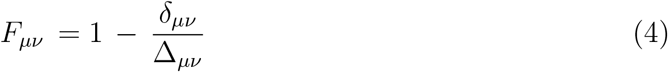

where *δ*_*µν*_ denotes the average distance between genomes *α* and *β* ≠ *α*, both in cluster *µ*, or both in cluster *ν*, and Δ_*µν*_ denotes the average distance between genomes *α* in cluster *µ*, and *β* in cluster *ν*. Accordingly, I refer to *δ*_*µν*_ and Δ_*µν*_ as the average within cluster and between cluster distances, respectively. Note that when genomes in clusters *µ* and *ν* are similar, *δ*_*µν*_ ≃ Δ_*µν*_ so that *F*_*µν*_ ≃ 0.

The simulations proceed as follows: A population of *N* identical genomes is first spread across the landscape at random. Initially, genomes contain no mutations (i.e., 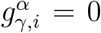 uniformly). In stage (i) of the algorithm, each individual undergoes a single random flight of mean length *λ* = 1, distributed as *f* (*x*) = *λ*^−1^ exp −*x/λ*. The boundaries of the landscape are reflective (see Online resource 1). In stage (ii), each female randomly selects a mate from among males within a distance *R* = *λ* of its position on the landscape, subject to the requirement that *q*^*αβ*^ ≥ *q*_*min*_ where *q*_*min*_ = 0.95. The number of offspring produced in a given mating event is Poisson distributed with mean

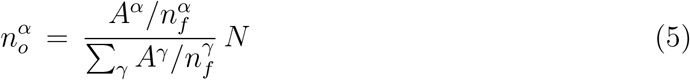

where *A*^*γ*^ is the area of overlap between the landscape and the mating circle for genome *γ*, 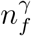 is the total number of female genomes within the mating circle (including *γ* itself), and the sum over *γ* in Eq. (5) is restricted to mated female genomes (see Online resource 1). As a result, the average total number of offspring produced is equal to *N*, and the number of offspring produced in a particular mating event decreases with the local density of females, reflecting the availability of resources in a natural population. Finally, in stage (iii), offspring genomes undergo random mutations, and are deposited at the positions of their respective female parents. The parent generation is removed, and the procedure starts again from stage (i).

Each of the simulations below run for 4*N* generations, and are sampled at intervals of Δ*t* = 100 generations. In all simulations, the mutation rate is *µ* = 0.05*/N* per gene per generation, slightly larger than the estimate above based on the effective population size for Heliconius. Populations evolve on landscapes of width *w* and length *l*. The mean number of individuals is *N* = *l w ρ*, where *ρ* is the mean number of individuals per unit area; Here, I consider the values *ρ* = 4, 8, and 16, consistent with density and flight length data available for non–migratory butterflies (this data is discussed in detail later below). I focus on the case *ρ* = 4 for which the average number of individuals in a mating circle, *πR*^2^*ρ*, is consistent with typical values simulated by de Aguiar et al. in [9–11]. Here and below, species clusters (Fig. 2) are computed from random samples of size min {*N/*4, 10^4^} individuals. To determine the number of species, I examine all partitionings up to 20 clusters. Genomes are partitioned using the “average” method defined in [27]. In all of the simulations, *F*^⋆^ = 0.24. The unit of length in the model is arbitrary, however, since *λ* = 1, all length parameters are in effect expressed in units of *λ*. For the species considered here, a typical value for *λ* would be in the range *λ* = 0.1 − 1 km. The data in this work were generated using a multi–thread algorithm [31] which divides the computational overhead of the simulations among 32 processor cores. The timescale for individual simulations ranges from hours to weeks depending on *N*.

**Fig. 2.**
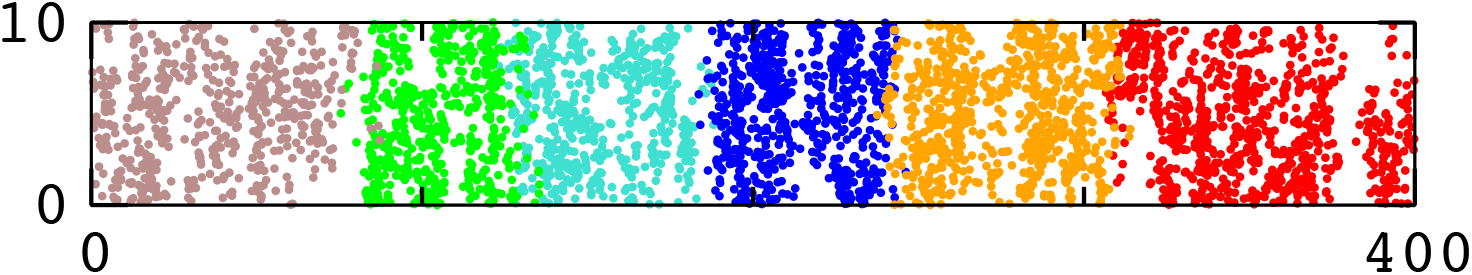
Snapshot of the speciation pattern taken during the simulation in Fig. 3 described later below.

## Results

To verify the model, I first examined some basic statistics along a trajectory. Fig. 3 shows the number of species, *ν*(*t*), as a function of time, *t* (where *t* is the number of generations) for a simulation with *ρ* = 4 and chromosome length *n* = 10^3^. In general, *ν*(*t*) reaches a steady state after about *N* generations for the parameters used in this work. Fig. 4 shows the relationship between *F*_*µν*_, Δ_*µν*_ and *δ*_*µν*_ for partitionings sampled during the simulation in Fig. 3; A fit of the function Δ = *δ /* (1 − *F*) obtained from Eq. (4) to the data for Δ_*µν*_ versus *F*_*µν*_ yields *δ ∼* 39 (correlation coefficient of 0.97). Notice that the smallest sampled distances between clusters have Δ_*µν*_ ∼ 50, in agreement with the value *d*_*max*_ = 50 used in the simulation (recall that *q*_*min*_ = 0.05).

**Fig. 3.**
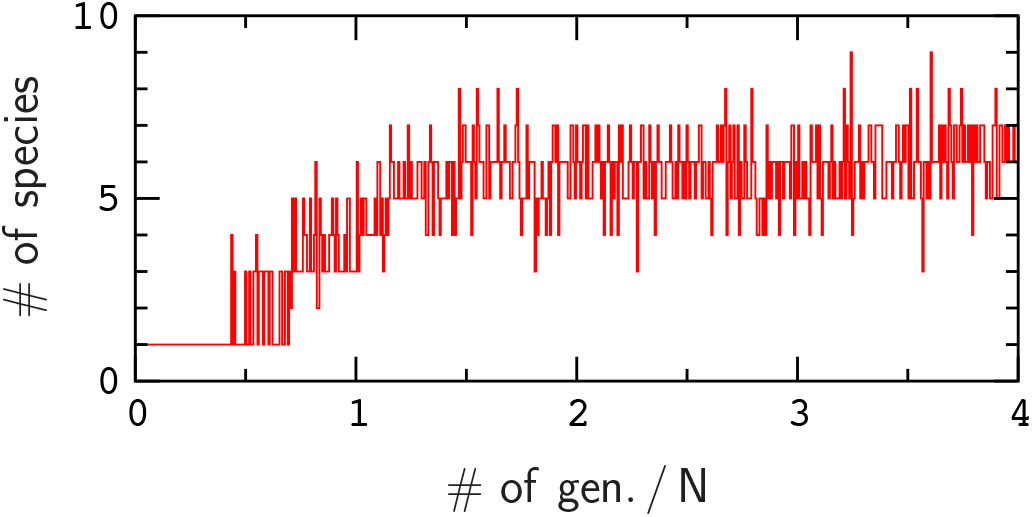
Number of species, *ν*(*t*), for a simulation with *ρ* = 4, *n* = 10^3^ and *F*^⋆^ = 0.24. The dimensions of the landscape are the same as in Fig. 2. *ν*(*t*) increases for smaller values of *F*^⋆^.

**Fig. 4.**
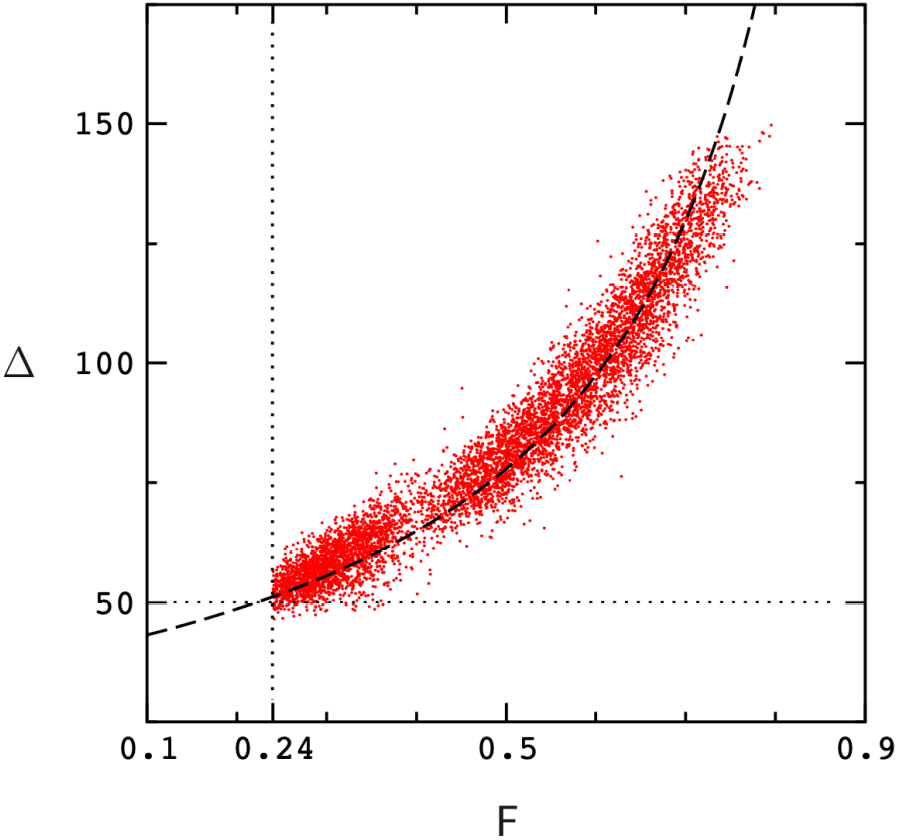
Correlation between Δ_*µν*_ and *F*_*µν*_ for partitions selected during the simulation in Fig. 3. The dashed line is a fit to the function Δ = *δ /* (1 − *F*) leading to *δ* ≃ 39. Dotted lines indicate the values of *F*^⋆^ and *d*_*max*_ used in the simulation.

In Fig. 5, I compute the local density of individuals during the simulation by overlaying the landscape with a square grid (with grid spacing *λ*) and then counting the number of individuals contained in each grid cell. Fig. 5 plots the probability distribution for the occupancy of grid cells. The distributions obey Poisson statistics except for small departures due to reflective boundary conditions (see Online resource 1). Fig. 6 shows the probability distribution for the number of mating events per male; The average number of mating events is 1, as expected for equal densities of male and female individuals.

**Fig. 5.**
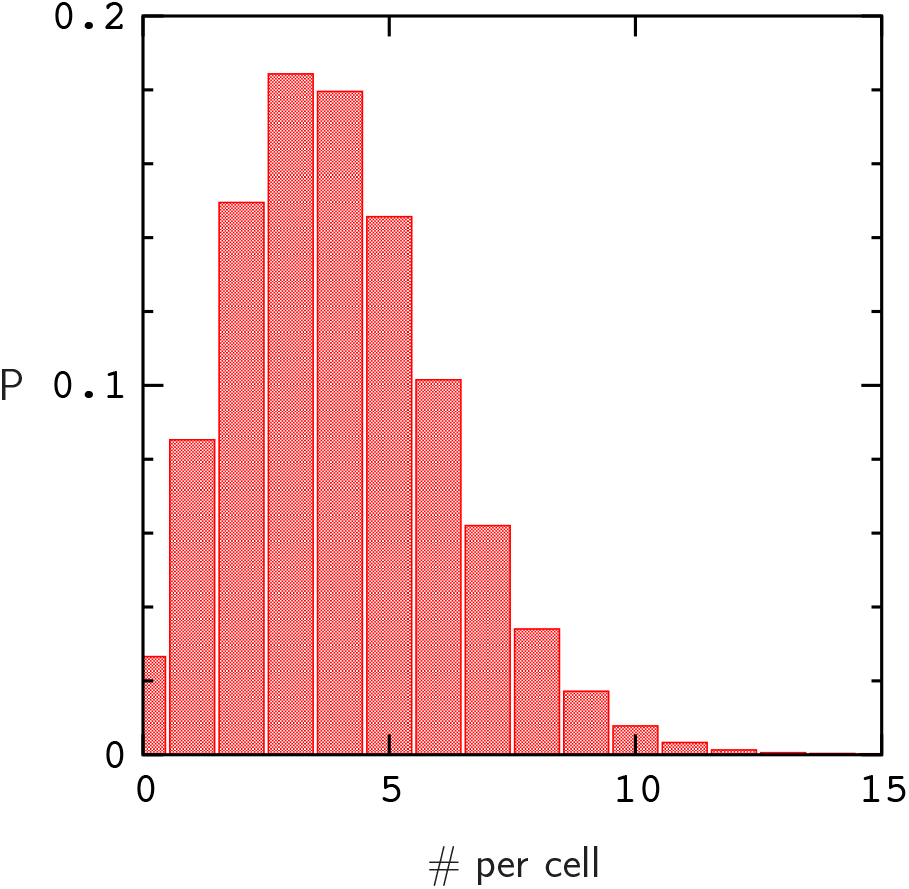
Probability distribution for the number of individuals per grid cell (see text) for the simulation in Fig. 3. The average number of individuals per cell is equal to *ρ*.

**Fig. 6.**
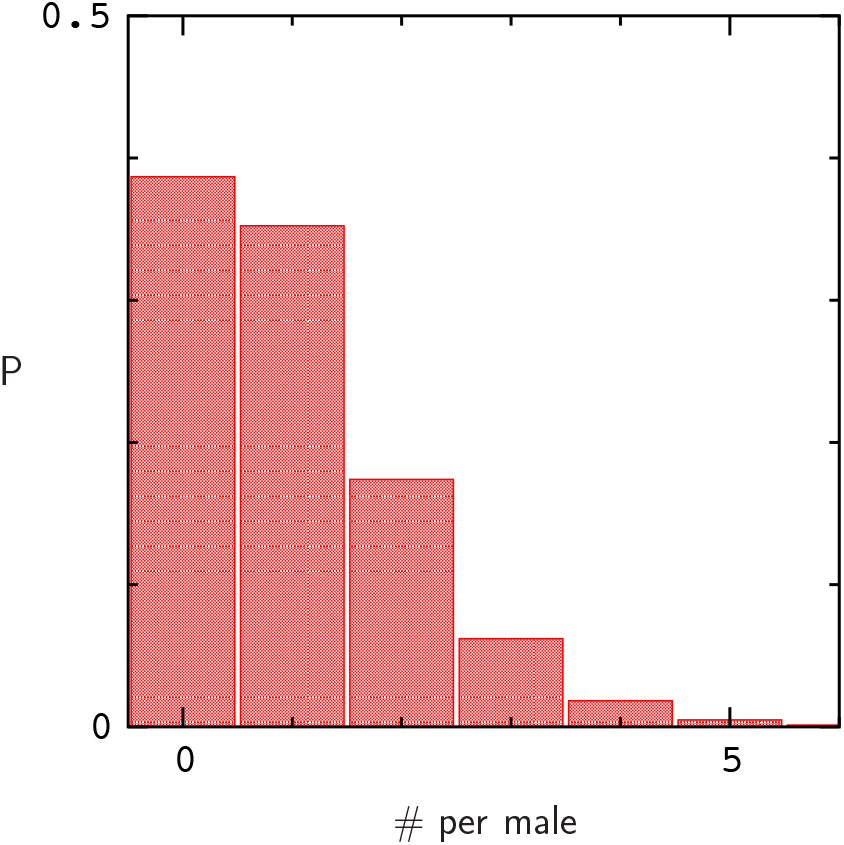
Probability distribution for the number of mating events per male individual for the simulation in Fig. 3.

The remaining figures describe speciation in the model as it is scaled toward real-istic values of *N* and *µ*. As noted above, this is accomplished by increasing the area of the landscape at constant *ρ* with *Nµ* = 0.05. Fig. 7 describes the average number of species in the steady state region, ⟨*ν*⟩_*t>N*_, for *ρ* = 4 on landscapes with fixed width and increasing length. Each data point corresponds to a single simulation of length 4*N*. In this case, ⟨*ν*⟩_*t>N*_ increases roughly linearly as the area of the landscape is increased, ultimately resulting in a narrow, almost 1–dimensional landscape as *µ* and *N* approach the values estimated for Heliconius “genes” via the condition *Nµ* = 0.05. By contrast, ⟨*ν*⟩_*t>N*_ approaches an asymptotic value with increasing area if the length to width ratio of the landscape is held constant (Fig. 8). In this case, simulations with relatively small population sizes roughly predict the behavior of ⟨*ν*⟩_*t>N*_ in much larger populations. An additional plot for *l/w* = 100*/*7 is provided in Online resource 1. The time scales of the simulations prevented an investigation of larger population densities.

**Fig. 7.**
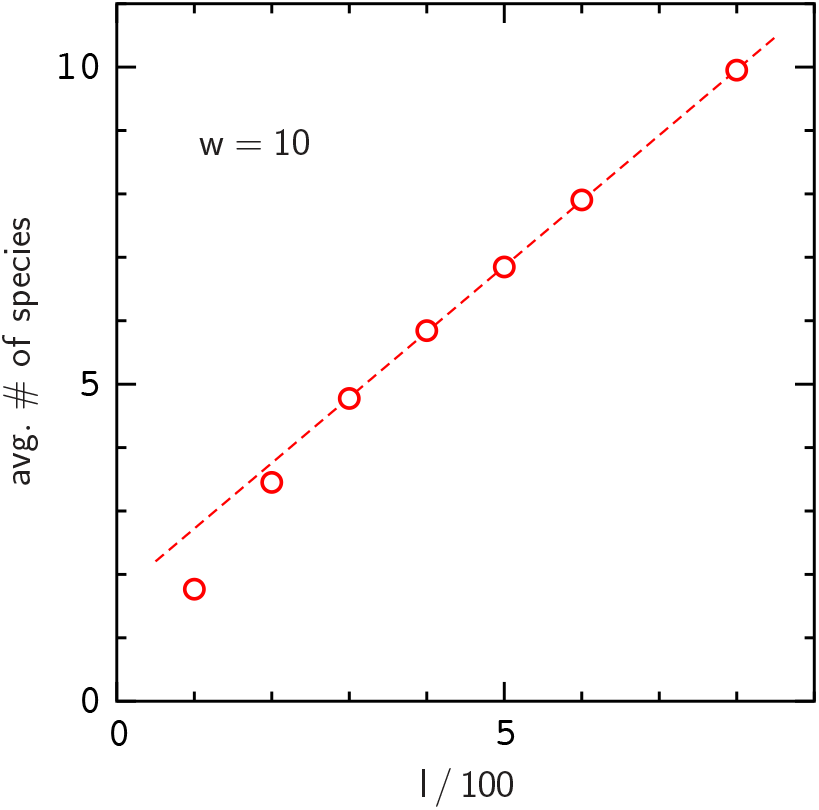
Number of species, ⟨*ν*⟩_*t>N*_, versus landscape length for simulations with *ρ* = 4, *n* = 10^3^, and *F*^⋆^ = 0.24. The dashed line is a fit to the data for *l/*100 *>* 2.

**Fig. 8.**
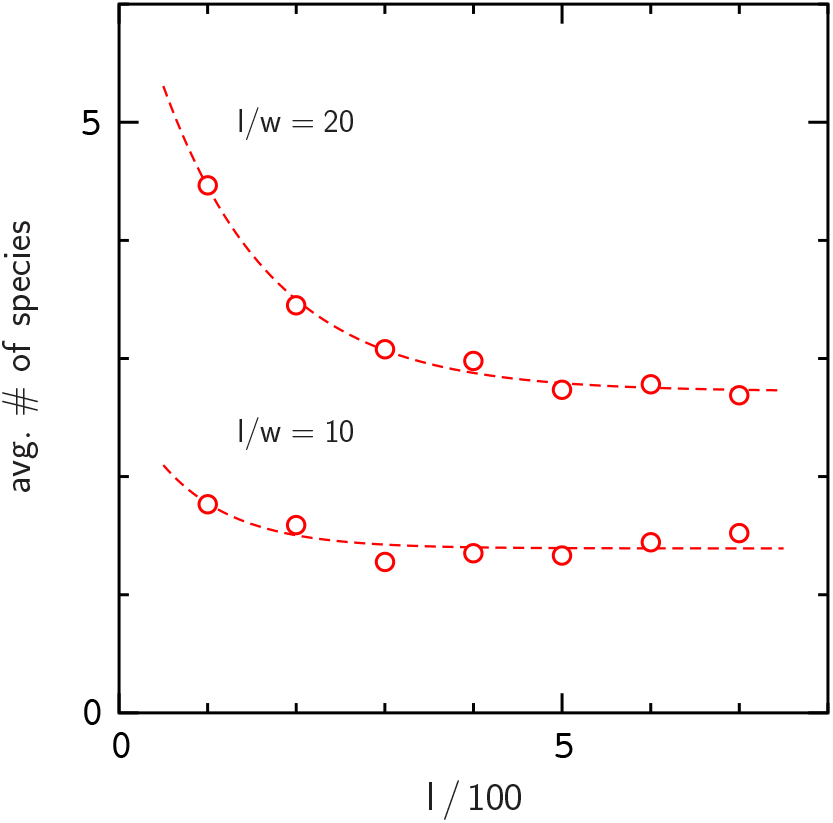
Number of species, ⟨*ν*⟩_*t>N*_, versus landscape length for landscapes with fixed length to width ratio. The parameters are *ρ* = 4, *n* = 10^3^, and *F*^⋆^ = 0.24. Dashed lines are fits of the function *f* (*l*) = *α* + *β* exp(−*γl*) to the data.

The results in Fig. 8 suggest that speciation will occur for *l/w* ⪞ 10. However, the data in [8] suggest that ⟨*ν*⟩_*t>N*_ is sensitive to genome length. To explore this possibility, I computed ⟨*ν*⟩_*t>N*_ for increasing genome lengths on fixed landscapes (Fig. 9). Note that the crossover rate remains constant regardless of genome length, as described above. Increasing the length of the genomes clearly seems to enhance speciation, however, this effect decreases significantly with increasing population density (see Online resource 1), and is generally smaller than expected from the theory given in [9].

**Fig. 9.**
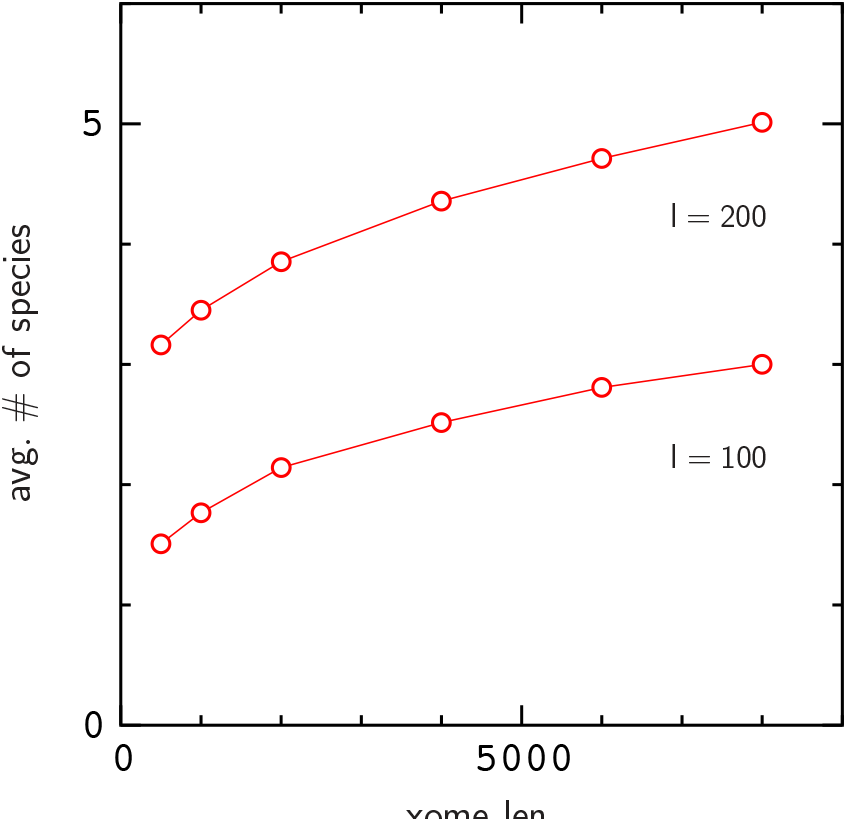
Number of species, ⟨*ν*⟩_*t>N*_, versus chromosome length for landscapes with fixed dimensions. In both plots, *w* = 10 and *ρ* = 4.

I also examined the dependence of ⟨*ν*⟩_*t>N*_ on population density. Fig. 10 describes the effect of increasing density on ⟨*ν*⟩_*t>N*_ for landscapes with fixed width *w* = 10 and increasing length (as in Fig. 7). For each landscape length, ⟨*ν*⟩_*t>N*_ decreases with increasing density. This result is consistent with the effect of increasing the mating radius (and, consequently, the number of individuals in the mating circle) in [11]. The effect of increasing the landscape width at higher densities is shown in Online resource 1.

**Fig. 10.**
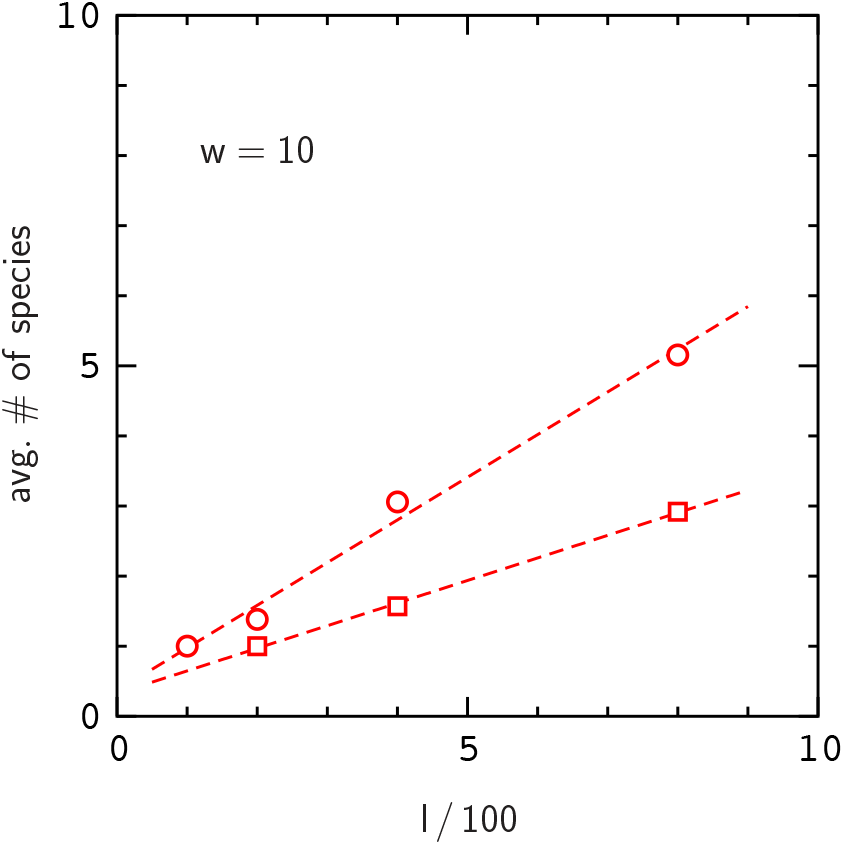
Number of species, ⟨*ν*⟩_*t>N*_, as in Fig. 7 for higher population densities, *ρ* = 8 (circles), and *ρ* = 16 (squares).

## Discussion

The results show that persistent species will appear in the model if it is continued to population sizes and mutation rates typical of butterflies; However, this depends on the dimensions of the landscape, and seems limited to landscapes with length to width ratio ⪞ 10, or perhaps ⪞ 5 depending on genome length, *F*^⋆^, and other parameters used in the simulations. The dependence of ⟨*ν*⟩_*t>N*_ on the length to width ratio of the landscape is consistent with the results in [7] where, for a given landscape area, species are much more abundant on 1–dimensional landscapes (i.e., with *l/w »* 1) than fully 2–dimensional landscapes with *l/w* = 1. A possible explanation for this effect is that persistent clusters are easier to maintain on narrow landscapes since they are exposed to gene flow from other species on only two small fronts (Fig. 2), whereas for *l/w* = 1 species clusters tend to be fully exposed [7, 8]. The dispersal mechanism used here also leads to more rapid gene flow across the landscape than the mechanism used by de Aguiar et al. Thus, given also the small value of *Nµ* used here, it is not difficult to see why speciation might be limited to narrow landscapes in the present model.

Aside from *Nµ*, the results depend on the limit *F*^⋆^, and the parameter *f* and *d*_*max*_ (which corresponds to the parameter G in [7]). It has been shown theoretically that the number of species in the de Aguiar model increases with both *N* and *µ* if other parameters such as population density and genome length are held constant [9]. Since our models are not too different, larger values of *Nµ* should lead to larger numbers of species in our model, while increasing *d*_*max*_ should lead to fewer species. In this work, I have used a conservative value for *F*^⋆^, however, a smaller value, such as *F*^⋆^ = 0.2, might also be considered reasonable [28], and would lead to a noticeable increase in ⟨ *ν* ⟩_*t>N*_ (but also to noisier trajectories for *ν*(*t*)). According to the data for butterflies, *f* is often larger than 0.01 for incipient species, which would lead to a lower estimate for *µ* via Eq. 2. It is also important to remark that natural resource distributions are often confined to narrow, or coastal regions, and are usually textured or fragmented in ways that can promote speciation [15, 32, 33]. Under these circumstances, speciation in the model could be more pronounced.

As noted earlier, the relationship between flight length and population density was constructed to agree with the parameters available for butterflies. While population density estimates are available for many types of butterfly, flight length data seems limited to threatened species; I was able to find both parameters for two species – the Fender’s blue and the Glanville fritillary. For the Fender’s blue, Schultz and Crone estimate the average length of a single flight within a resource patch at 2.1 m, with about 2100 such flights per lifetime [25]. Assuming a random walk, this leads to a per generation flight length of about *λ* = 0.1 km. From the data available in [25], the typical number of individuals within an area of size *λ*^2^ is between 3–5. For the Glanville fritillary [34], I obtain a rough estimate for the per generation flight length of about *λ* = 1 km, and a population density of between 2–12 individuals per km^2^ [35]. The density values used in the model (in particular, the value *ρ* = 4) are consistent with these estimates.

Finally, while the data in Fig. 8 clearly indicate scaling of species numbers with *Nµ*, more data is needed to establish this result over a wider range of population sizes and densities. Unfortunately, despite parallelization of the problem, a study of this kind would be very difficult with the present algorithm (for example, the largest populations in Fig. 8 (*N* ≃ 2.5 *×* 10^5^) required over three weeks of simulation time – a requirement that grows linearly with population size). More realistic population sizes of order 10^6^ or larger should be tractable on graphical processing units, where the number of available processor cores is orders of magnitude larger [36]. This sort of approach could provide a much clearer picture of neutral speciation in natural populations.

## Supporting information

Online resource 1

## Acknowedgments

I thank Nick Grishin for providing access to computer facilities during this work.

## Data Availability

Data for this work is contained in the manuscript file.

