## Supplementary material for "Neutral speciation in realistic populations": Online resource 1

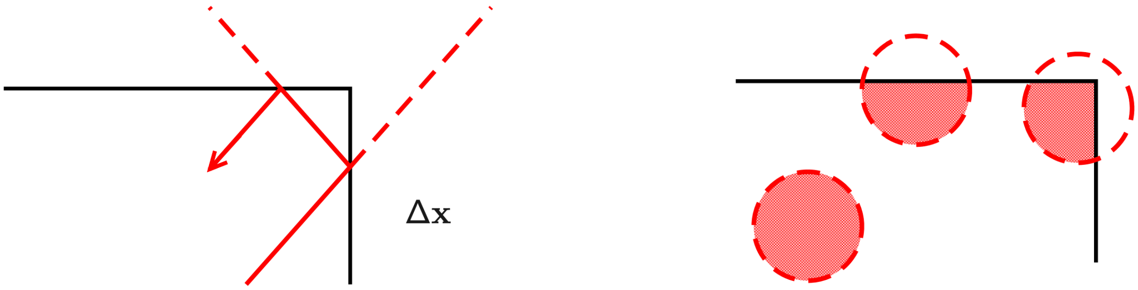


**Fig. 1** Landscape boundaries are reflective: Flights $\mathbf{\Delta x}$ (left panel) that would extend outside the landscape (as indicated by dashed lines) are continually reflected until they terminate within the landscape. The area of overlap $A^{\gamma}$ between the mating circle and the landscape for a focal individual $\gamma$ depends on its position on the landscape (right panel). $A^{\gamma}$ is determined analytically in the simulations.

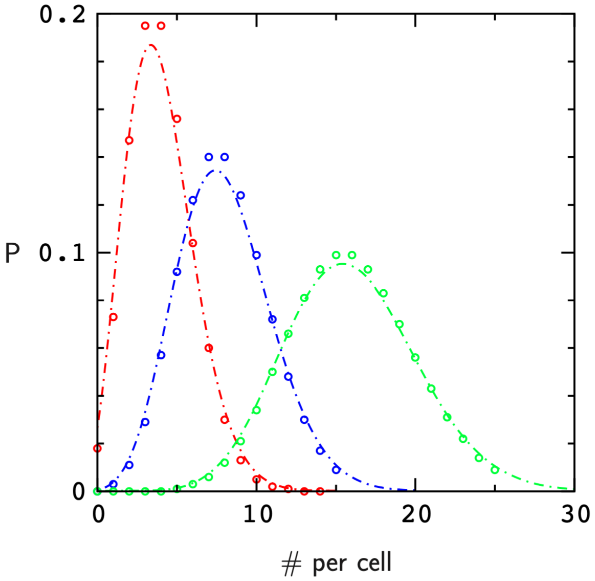


**Fig. 2** Probability distributions for the number of individuals per grid cell (dashed lines) for $\rho = 4$ (red), $\rho = 8$ (blue) and $\rho = 16$ (green) and predicted values (circles) based on the Poisson distribution, $P(k) = \rho^k \exp(-\rho)/k!$, where $k$ is the number per cell. The average number of individuals per cell in each simulation is $\rho$ (the landscape dimensions are $l = 400$ and $w = 10$). Small departures from Poisson statistics are likely due to reflective boundary conditions: Each individual produces the same number of offspring (on average) per generation. Near the edges of the landscape, the density is a little larger since some of the flight paths are reflected back to the interior at the start of each generation.

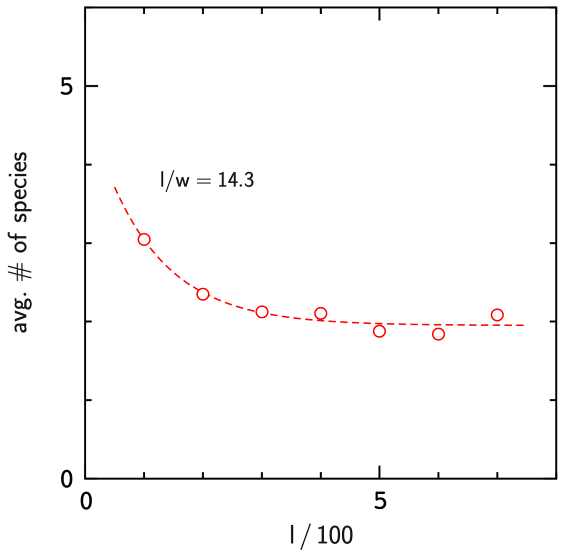


**Fig. 3** Number of species, $\langle\nu\rangle_{t>N}$, versus landscape length for a landscape with length to width ratio $l/w = 100/7$. The parameters are $\rho = 4$, $l = 10^3$, and $F^\star = 0.24$. The dashed line is a fit of the function $f(l) = \alpha + \beta\,\exp(-\gamma l)$ to the data, as in Fig. 8 of the Text.

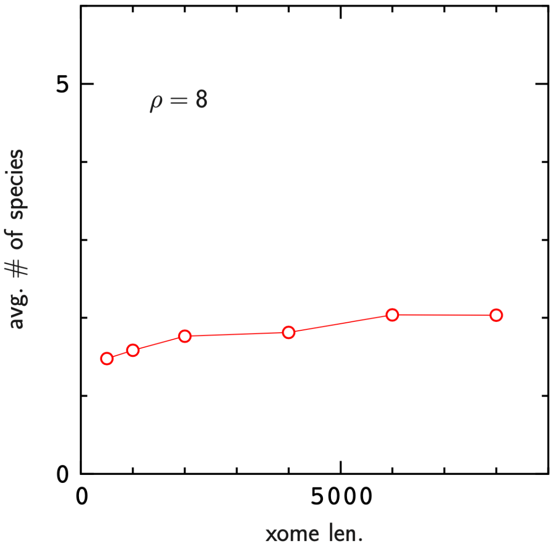


**Fig. 4** Number of species, $\langle \nu \rangle_{t>N}$, versus chromosome length as in Fig. 9 of the Text. The parameters are $\rho = 8$, $w = 10$, and $l/w = 20$.

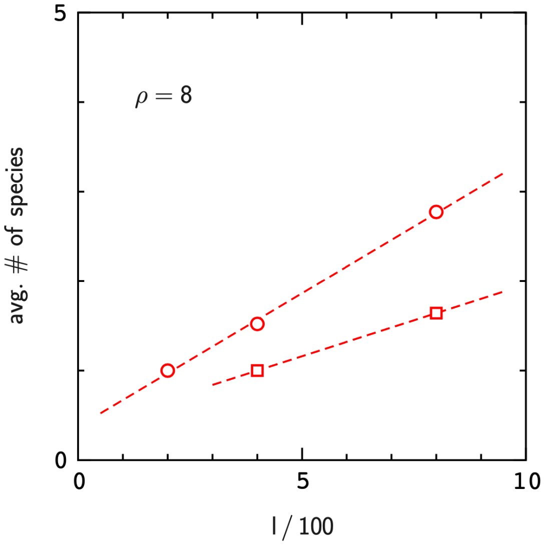


**Fig. 5** Number of species, $\langle \nu \rangle_{t>N}$, as in Fig. 9 for wider landscapes, $w = 20$ (circles) and $w = 40$ (squares).
